# *Alk1* mutant endothelial cells undergo clonal expansion in mouse brain arteriovenous malformations

**DOI:** 10.1101/2020.05.20.106799

**Authors:** Sonali S. Shaligram, Rui Zhang, Wan Zhu, Li Ma, Ethan Winkler, Man Luo, Qian Li, Thomas Arnold, Grez NG Santander, Cameron M. McDougall, Julia Wong, Rich Liang, Leandro Barbosa Do Prado, Chaoliang Tang, Hua Su

## Abstract

**Rationale:** Mutation in human arteriovenous malformation (AVM) causative genes in a fraction of endothelial cells (ECs) causes AVMs in mice. It is unclear how a small number of mutant ECs can lead to AVM formation.

**Objective:** To understand how a fraction of mutant ECs causes AVM, we tested the following hypotheses: (1) activin receptor-like kinase 1 (*Alk1* or *Acvlr1*) mutant brain ECs undergo clonal expansion upon angiogenic stimulation, (2) *Alk1* mutant ECs display growth advantage, (3) the burden of *Alk*1 mutant ECs correlates with AVM severity, and (4) *Alk1* mutant bone marrow (BM) derived ECs alone is sufficient to cause AVM.

**Methods and Results:** We used *Pdgfb*iCreER;*Alk1*^f/f^;confetti^+/−^ mice which express an EC-specific tamoxifen (TM)-inducible Cre recombinase, a Cre-regulated confetti transgene, and *Alk1* floxed alleles. Brain AVMs were induced by direct brain injection of an adeno-associated viral vector expressing vascular endothelial growth factor (AAV-VEGF) followed with intra-peritoneal injection of TM two weeks later. Color-predominance of confetti reporter in AVMs compared to control brain ECs suggested that clonal expansion was associated with AVM development. We treated *Pdgfb*iCreER;*Alk1*^f/f^ with different doses of TM to create a mosaic of wild-type (WT) and mutant ECs and found that equal numbers of Alk1^+^ and Alk1^−^ ECs were proliferating. Increase of TM dose increased the number of Alk1^−^ ECs, the abnormal vessels in brain AVMs, the number of arteriovenous shunts in the intestines, and mouse mortality. To test if mutation of Alk1 in BM-derived ECs can cause brain AVM, we transplanted WT mice with BM of *Pdgfb*iCreER;*Alk1*^f/f^ mice. After AAV-VEGF and TM treatment, these mice developed AVMs in their brains and arteriovenous shunts in their intestines.

**Conclusion:** Clonal expansion of *Alk1* mutant ECs could partly explain why a fraction of mutant ECs causes AVM. Mutation of AVM causal genes in BM-derived ECs is sufficient to cause AVM formation.

## INTRODUCTION

Arteriovenous malformations (AVMs) are tangles of abnormal vessels that shunt blood directly from arteries to veins. Vessels in AVM have abnormal wall structure and are prone to rupture causing life-threatening hemorrhage [1]. Most AVM cases are sporadic. There are a few familial cases, such as those with hereditary hemorrhagic telangiectasia (HHT), an autosomal dominant disorder. HHT patients have a high prevalence of AVMs in multiple organs including the brain, lung, and intestine [1, 2]. Although HHT causal genes are mostly identified, such as, endoglin (*ENG*), activin receptor-like kinase 1 (*ALK1* also known as *ACVLR1*), and *SMAD4* [1, 2], it is still not clear how AVMs are formed.

The familial forms of the more common sporadic disorders have been used to study the disease mechanisms of sporadic cerebrovascular diseases [3, 4]. We have established several AVM mouse models that have the AVMs in multiple organs and in the brain angiogenic region through conditional deletion of the HHT causative gene, *Eng* or *Alk1* in adult mice, combined with brain focal angiogenic stimulation [5–7]. Using these models, we found that deletion of *Alk1* or *Eng* in endothelial cells (ECs) and angiogenic stimulation are necessary for AVM formation in the brain of adult mouse [6–8]. Deletion of *Alk1* or *Eng* in ECs is necessary for AVM formation in other organs too [9]. We also found that homozygous mutation of *Alk1* or *Eng* in a fraction of somatic ECs [5, 10] is sufficient to induce AVM phenotype in the brain angiogenic regions and in the intestines of adult mice. In addition, we showed that transplantation of *Eng*-deficient bone marrow (BM) cells caused cerebrovascular dysplasia in wild-type (WT) mice after VEGF stimulation [11], suggesting a potential contribution of BM-derived cells in the pathogenesis of AVM. Interestingly, recent studies showed that sporadic brain AVMs and extra-neural AVMs harbor somatic mutations of genes in Ras-mitogen-activated protein kinases (MAPK) pathways including KRAS, MAP2K1 and BRAF. These mutations were mostly present in ECs [12–14]. However, it is not clear how a fraction of mutant ECs leads to AVM development and if mutation in BM-derived ECs alone can cause AVM formation.

EC *Eng* deletion in neonates increases proliferation of both *Eng*-deleted and WT ECs in AVM vessels [15]. We found that *Alk1*-deleted ECs preferentially cluster in brain AVM vessels [7]. As it is statistically unlikely that the dominance of recombined cells in AVMs is the result of multiple independent Cre recombination events, the observed pattern indicates that mutant ECs may undergo clonal expansion. To determine if mutant ECs are undergoing clonal expansion, we used used R26R-confetti reporter [16] to track progression of individual recombined cells in mice with *Alk1* EC mutation and brain AVMs. Confetti transgene contains two loxP-flanked dimers; one dimer has nuclear-localized green green fluorescent protein (GFP) and reversed cytoplasmic yellow fluorescent protein (YFP); the other dimer has cytoplasmic red fluorescent protein (RFP) and a reversed membrane-tethered cyan fluorescent proten (CFP). R26R-confetti reporter mice were crossbred with *Pdgfb*iCreER;*Alk1*^f/f^ mice which express an EC-specific tamoxifen (TM)-inducible Cre recombinase, and *Alk1* floxed alleles. Upon TM treatment, Cre will be expressed in ECs, triggering *Alk1* deletion in the ECs, at the same time activate the expression of 1 of the 4 confetti fluorescent genes in individual ECs. The progenies derived from single EC will express same fluorescent protein. Thus, this reporter provides a unique way to identify a cluster of cells that are derived from a single parental cell-indicating clonal expansion [17, 18].

In this paper, using *Alk1* conditional knockout adult mice, we showed that *Alk1* mutant ECs undergo clonal expansion in brain AVMs. The number of Alk1^−^ ECs is positively correlated with AVM phenotype severity. Mutation of *Alk1* gene in BM-derived ECs alone is sufficient to cause AVM development.

## MATERIALS AND METHODS

### Animals

The protocol and experimental procedures for using laboratory animals were approved by the Institution of Animal Care and Use Committee (IACUC) at the University of California, San Francisco (UCSF). Animal husbandry was provided by the staff of the IACUC, under the guidance of supervisors who are certified Animal Technologists, and by the staff of the Animal Core Facility. Veterinary care was provided by IACUC faculties and veterinary residents located on the Zuckerberg San Francisco General Hospital campus.

*Pdgfb*iCreER;*Alk1*^f/f^;confetti^+/−^ mice that express an EC-specific TM inducible Cre recombinase, have their *Alk1* gene exons 4-6 floxed [19], and carry the R26R-confetti Cre-regulated transgene were used to test the clonal expansion of *Alk1* mutant ECs. R26R-confetti transgene (*Gt(ROSA)26Sortm1(CAG-Brainbow2.1)Cle*/J, the Jackson laboratory, Bar Harbor, ME) has a loxP-flanked STOP cassette between the CAG promoter and the Confetti sequence. Upon TM treatment, cells carrying Confetti locus will express 1 of the 4 confetti colors (cytoplasm RFP or YFP, nuclear GFP or membrane CFP), which is stably propagated within progeny [16]. The *Pdgfb*iCreER;*Alk1*^f/f^;confetti^+/−^ mice were treated with a single i.p. injection of 2.5 mg/25g TM.

The *Pdgfb*iCreER;*Alk1*^f/f^;Ai14^+/−^ mice that carry an Ai14 reporter were treated with different doses of TM through single i.p. injections to induce different levels of *Alk1* gene deletion and Ai14 expression in ECs. *Pdgfb*iCreER;*Alk1*^f/f^;Ai14^+/−^ mice were also used as BM donors and other experiments.

### Statistical Analysis

All quantifications were done by at least two researchers who were blinded to the treatment groups on images that were coded. Data were analyzed using GraphPad Prism 6 by one-way ANOVA followed with multiple comparisons and Tukey’s correction. Survival rate was analyzed using log-rank test. Data are presented as mean ± standard diviation (SD). A *P* value ≤0.05 was considered significant. Sample sizes were indicated in the figure legends.

The corresponding author will make all data, methods used in the analysis, and materials used to conduct the experiments available upon request. This material will be available at University of California, San Francisco, California.

Please see Online Data Supplement of Detailed Methods.

## RESULTS

### Alk1^−^ ECs undergo clonal expansion

Since we found previously that mutation of *Eng* or *Alk1* gene in a fraction of ECs is sufficient to cause AVM in the brain [5, 6] and *Alk1*-deleted ECs preferentially cluster in brain AVM vessels [7], we hypothesized that individual mutant ECs proliferate and clonally expand to cause AVM. Brain AVMs were induced in *PdgfbiCreER*;*Alk1*^2f/2f^;confetti^+/−^ mice through intra-brain injection of an adeno-associated viral vector expressing vascular endothelial growth factor (AAV-VEGF, 1×10^9^ gcs) to induce brain focal angiogenesis followed by i.p. injection of TM (2.5 mg/25g of body weight) 2 weeks later (Fig. 1A and Supplementary Fig. 1). TM treatment will induce *Alk1* gene deletion and confetti locus activation in ECs. The confetti reporter irreversibly labels individual recombined ECs with one of four colors. Therefore, if a single cell (of a single color) clonally expands, we would expect to see predominance of individual colors and closer proximity of individual recombined cells of a single color, that would occur at random. Indead, through observed brain sections under confocal microscopy 8 days after TM treatment, we found clusters of ECs-labelled with same confetti color in the brain AVMs, which were not detected in the brain with *Alk1* deletion alone or brain angiogenic region of wild-type (WT) mouse (Fig. 1B). The overall recombination rate and proportion of all colors were higher in brain AVMs than in WT brain angiogenic region and in brain with *Alk1* mutation in the ECs alone (Fig. 1C). Different proportion of confetti color labeled ECs were observed on brain AVMs of different mice and on different sections of same brain AVM (Fig. 1D). These data indicate that *Alk1*-mutant ECs are undergoing clonal expansion in brain AVM lesions.

**Fig. 1:**
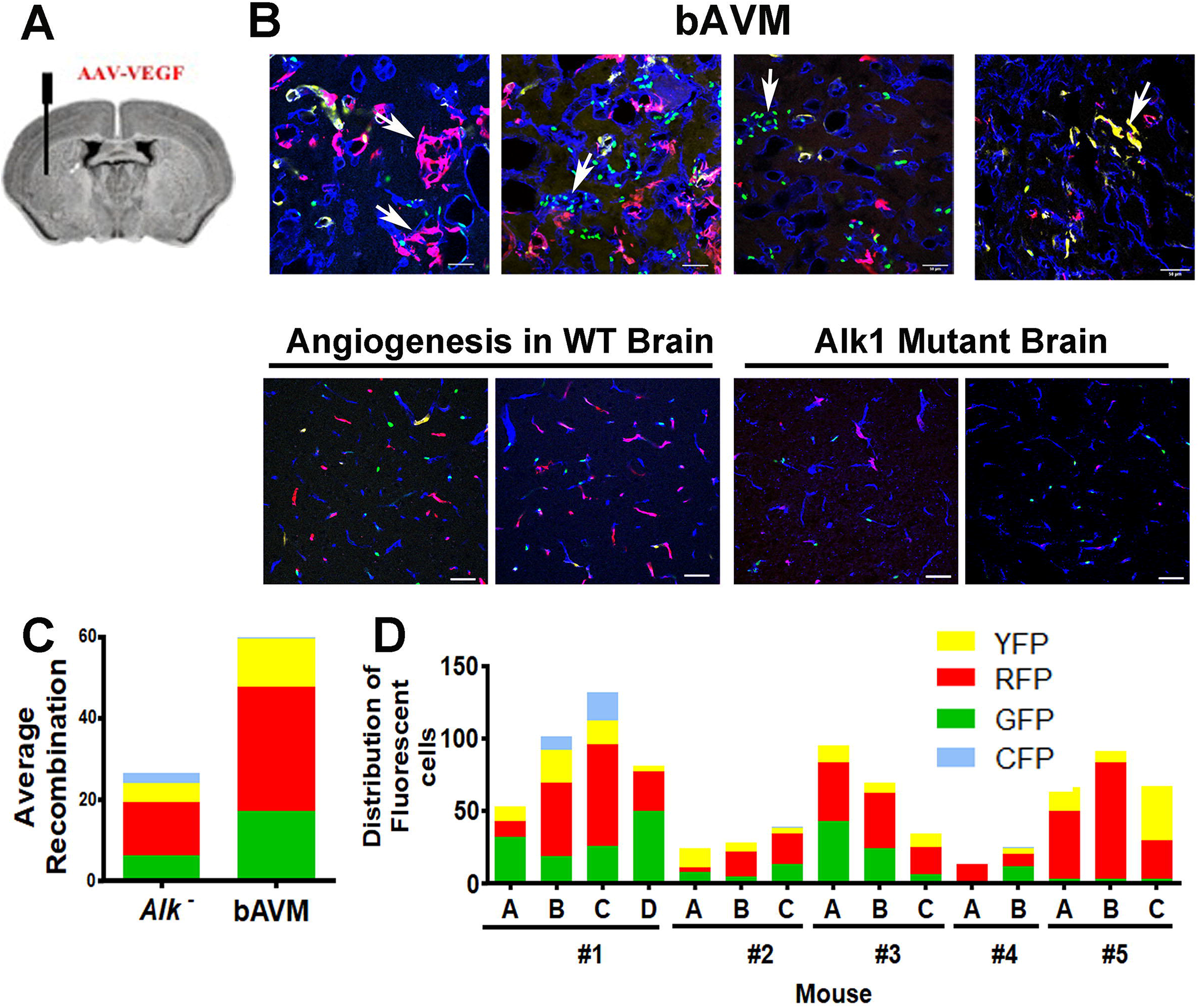
Clonal Expansion of ECs in brain AVMs. **A.** A image of brain coronal section illustrates the AAV-VEGF injection site during model induction. The AVM lesion will develop around the tip of the needle. **B.** Confocal images taken around the needle injection sites shown in **A**. Upper panel: brain AVM lesions. ECs were stained with an anti-CD31 antibody (blue). ECs expressing same confetti color were clustered together (Arrows). Bottom panel: confocal images of angiogenic region in WT brain (left) and confocal images of brains with *Alk1* mutation in ECs (right). ECs expressing individual confetti colors were scattered in these brain sections. **C**. Average number of ECs expressing confetti colors in 5 brain AVMs and 5 brains with *Alk1* mutatin in ECs alone. **D**. Distribution of confetti colors in each sections (A, B, C or D) analyzed in brain AVM lesions of 5 mice *(#1* to *#5*). Different proportions of confetti color labeled ECs were observed on different sections. Scale Bars=50 μm.

### Equal numbers of Alk1^+^ and Alk1^−^ ECs were proliferating in brain AVM lesions

We showed previously that more ECs were proliferating in brain AVM lesion that have *Alk1* deletion in near 100% ECs than in brain angiogenic region of WT mice [7]. But information on potential cell-autonomous effects in AVM development is lacking. To answer this question, we tested if reduce TM dose could reduce recombination in ECs to create brain AVM lesions with mosaic Alk1^+^ and Alk1^−^ ECs. AVMs were initiated by injecting AAV-VEGF into *Pdgfb*iCreER;*Alk1*^f/f^;Ai14^+/−^ mice, and gene recombination was induced by treating the mice with an high dose of TM (1.25 mg/25 g of body weight) and a low dose of (0.01 mg/25 g of body weight) through a single i.p. injection, Supplementary Fig. 1B). Brains were collected at 22 or 28 days after model induction, sectioned and stained for Alk1 (to assess degree of gene knockout), lectin or CD31 (to mark all ECs) and Ai14 (which marks recombined cells). The degree of recombination was calculated as a fraction of total ECs. Alk1 expression was calculated by immunofluorescence and co-localization with ECs and was verified by western blot to quantitatively assess whole brain Alk1 protein levels in mice treated with different TM doses. As expected, higher dose TM resulted in a greater degree of recombination and Alk1 mutation (based on absence of Alk1 immunostain). Mice treated with 1.25 mg/25g body weight of TM, have more Ai14^+^ ECs than mice treated with 0.01 mg/25g of TM in the brain AVM lesions (0.01 TM group, 61±20%, 1.25 TM group: 87%±5%, P=0.01, Fig. 2A and B) and in intestines (0.01 TM group, 38±9, 1.25 TM group: 60±12, P=0.019, Supplementary Fig 2). The TM vehicle (corn oil) treated mice also showed a low (but not zero) degree of recombination (29%±10%), possibly due to Cre leakiness (Fig. 2A and B). Increase TM have also decreased the number of Alk1^+^ ECs in brain AVM lesions (Control: 99±8.25%, 0.01 TM group: 48.01±16.8%, 1.25 TM: 2.3±4.34%, P<0.001, Fig. 2C & D) and Alk1 expression in the brains (Fig. 3).

**Fig. 2:**
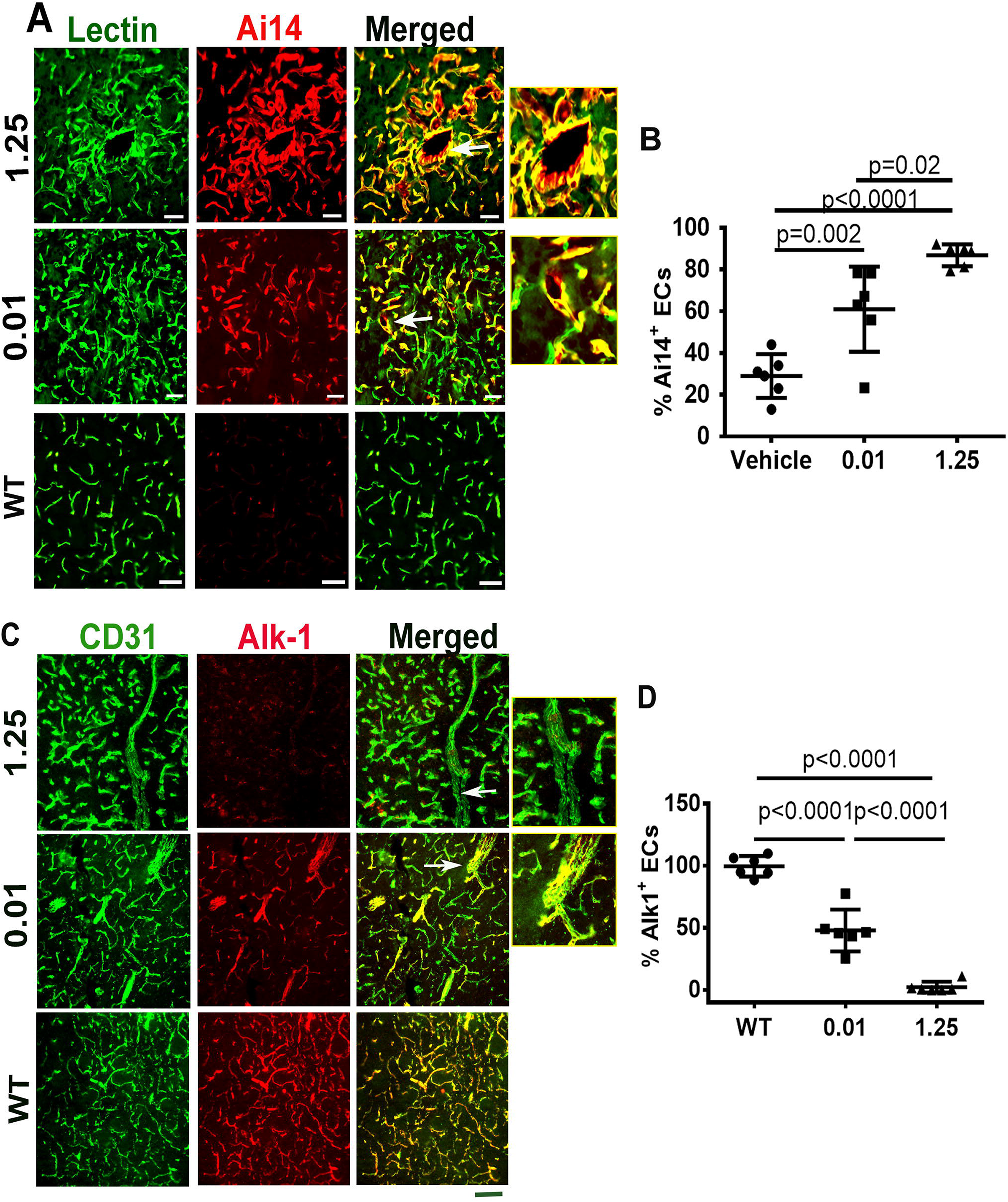
Increase of TM dose increased Ai14^+^ and reduced Alk1^+^ ECs. **A.** Representative images of sections collected from lectin (green) perfused brains. Cre activation in ECs was indicated by the expression of Ai14 reporter gene (red). Arrows indicate dysplastic vessels. The right side pictures show enlarged images of abnormal vessels indicated by arrows in the left side pictures. **B.** Quantification of Ai14^+^ ECs in brain AVM lesions. **C.** Representative images of brain sections co-stained with anti-CD31 (green) and anti-Alk1 (red) antibodies. Arrows indicate abnormal vessels. The right side pictures show enlarged images of abnormal vessels indicated by arrows in the left side pictures. **D.** Quantification of Alk1^+^ ECs. WT: corn oil treated mice; 0.01 or 1.25: mice treated with 0.01 mg/25g or 1.25 mg/25g TM. Scale bars=50 μm. N=6.

**Fig. 3:**
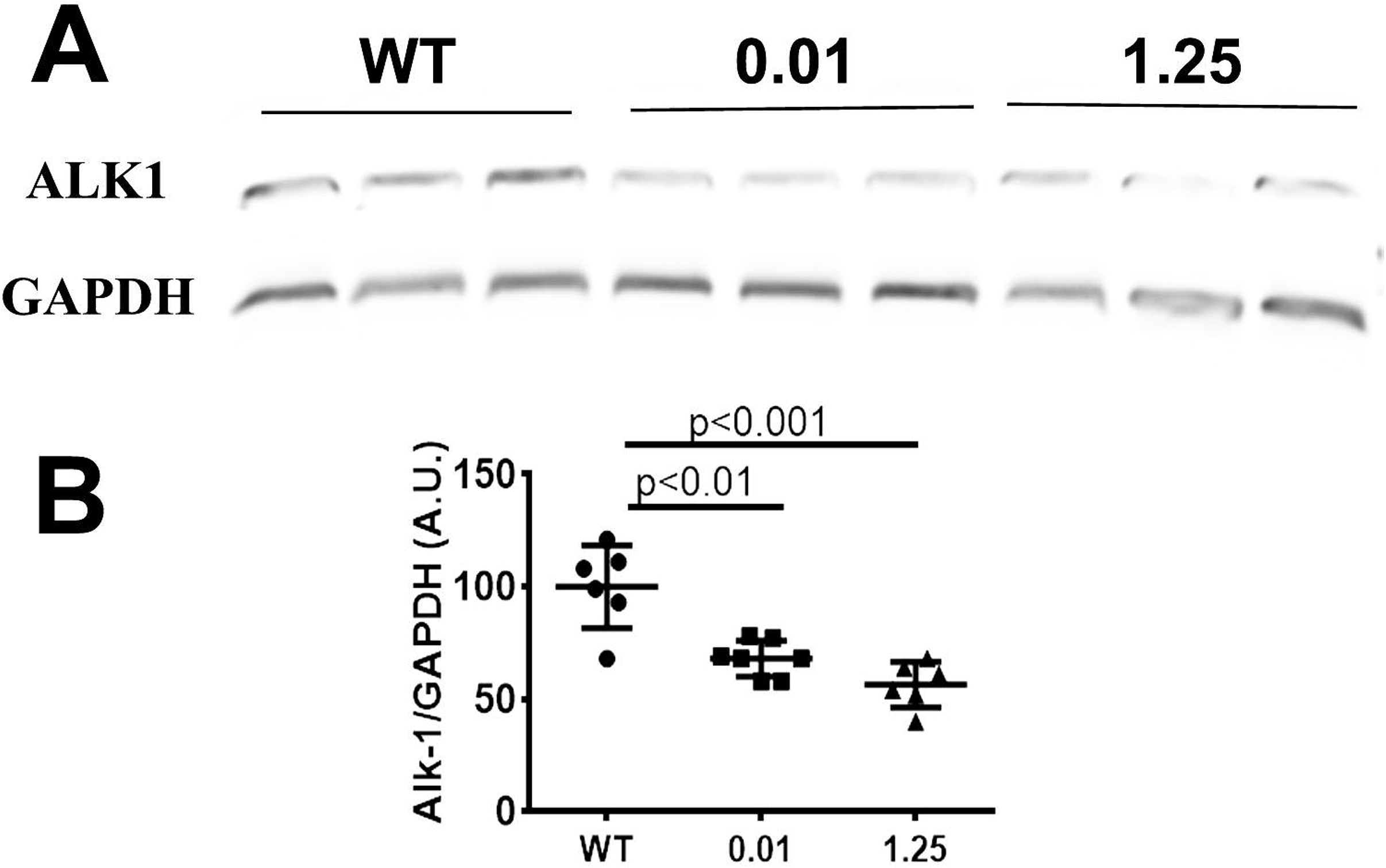
Increase of TM dose decreased Alk1 expression in brain AVM lesion. **A.** Representative western blot images. **B.** Quantification of Alk1 expression. WT: corn oil treated mice; 0.01 or 1.25: mice treated with 0.01 mg/25g or 1.25mg/25g TM. N=6 for WT and 1.25 group; N=7 for 0.01 group. A.U.: arbitrary unit.

We then analyzed EC-proliferation in brain AVM lesions with mosaic Alk1^+^ and Alk1^−^ ECs in mice treated with low-dose (0.01 mg/25g of body weight) TM. We co-labeled brain tissue sections using the proliferation marker Ki67, Erg (an pan-EC transcription factor), and Alk1 (to identify Alk1^−^ and Alk1^+^ ECs within mosaic lesions). We observed that 45% of Ki67^+^ ECs were Alk1^−^ ECs and 55% Ki67^+^ ECs were Alk1^+^ ECs (p=0.423, Fig. 4), indicated that Alk1^+^ and Alk1^−^ ECs have similar proliferation capacity in brain AVM lesions.

**Fig 4:**
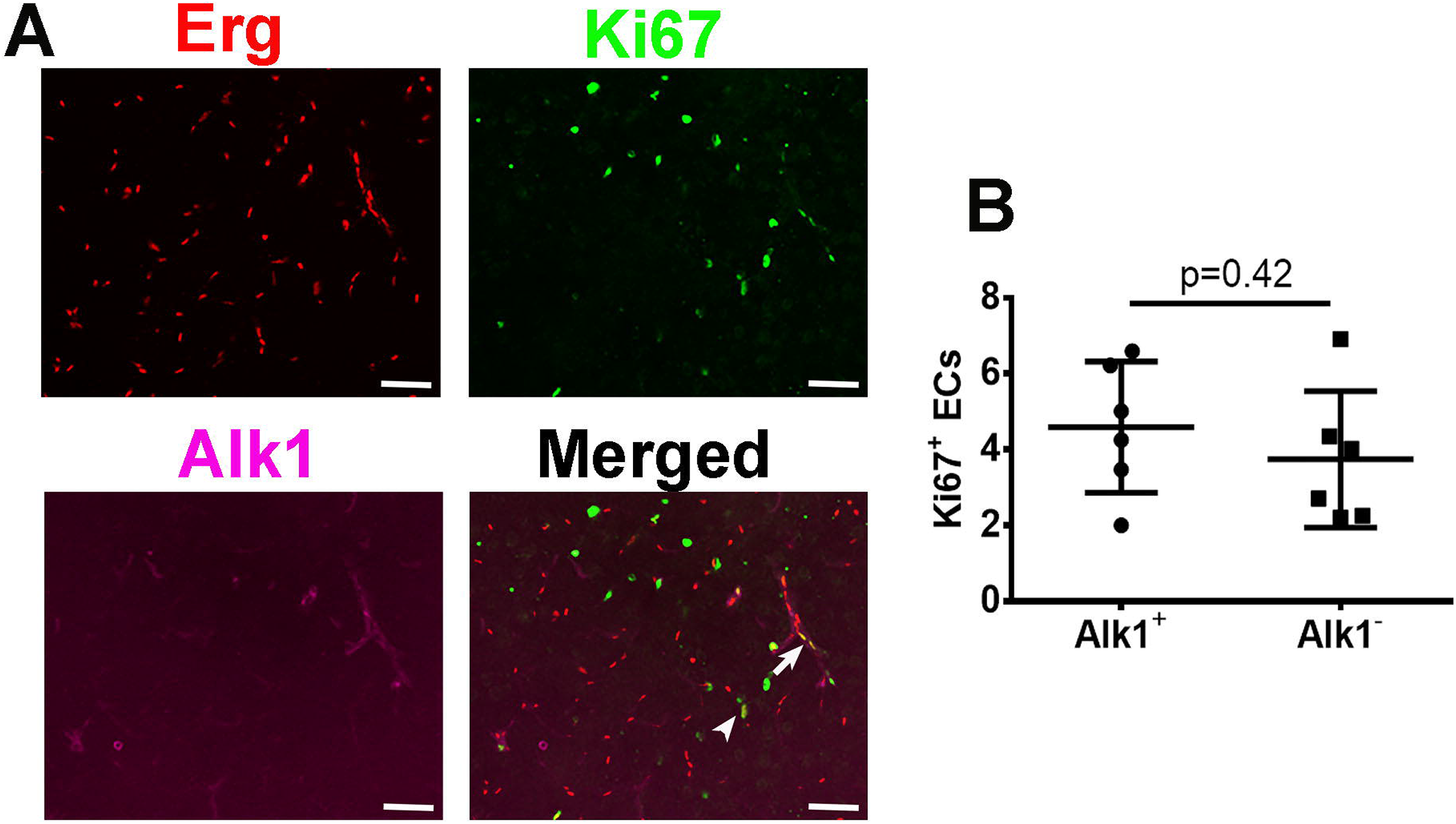
Equal number of Alk1^+^ and Alk1^−^ ECs were proliferating in brain AVM lesions. **A.** Representative images of brain AVM lessions. The nuclei of ECs were stained by an Erg specific antibody (red). Proliferating cells were stained by an Ki67 specific antibody (green). Alk1 expression was visualized by staining with an Alk1 specific antibody (purple). Arrow indicates a Ki67^+^ Alk1^+^ EC. Arrow head indicates a Ki67^+^ Alk1^−^ EC. Scale bars=50 μm. **B.** Quantification of Ki67^+^ Alk1^−^ and Alk1^+^ ECs. N=6.

### ALK1 mutation burden in ECs correlates with AVM severity and mouse mortality

Based on our data above, which suggested both cell-autonomous (clonal expansion) and non-cell autonomous (cell proliferation) functions in AVM, we were interested to understand the relationship between mutant EC burden and AVM severity.

To determine whether there is an association between number of mutant ECs and AVM severity, we quantified dysplastic vessels (expressed as dysplasia index: number of vessels with lumens >15 mm per 200 vessels) in brain AVM lesions and counted the number of intestinal arteriovenous (AV) shunts using latex dye casting. We found that mice in 0.01 TM group (4.5±1.1) have fewer dysplastic vessels in the brain AVMs than mice treated in 1.25 TM group (7.9±1.2, P=0.001, Fig. 5A and B). No dysplastic vessels were found in vehicle treated mice. Vessel densities were similar among groups (Fig. 5C). 100% of 1.25 TM treated mice had macroscopic level of brain AVM lesions, while only 50% of 0.01 TM treated mice developed macroscopic level of brain AVMs. The phenotype of 0.01 TM treated mice were analyzed 28 days after intra-brain injection of AAV-VEGF (Supplementary Fig. 1), which is 6 days later than 1.25 TM treated mice. No macroscopic level of AVMs was detected in vehicle treated mice (Supplementary Fig. 3A). Reduction of TM dose also reduced AV shunts in the intestines (Supplementary Fig. 3B, and Supplementary Table 1) and mouse mortality (Fig. 5D). Together, these data indicated that the burden of mutant ECs correlates with AVM severity.

**Fig. 5:**
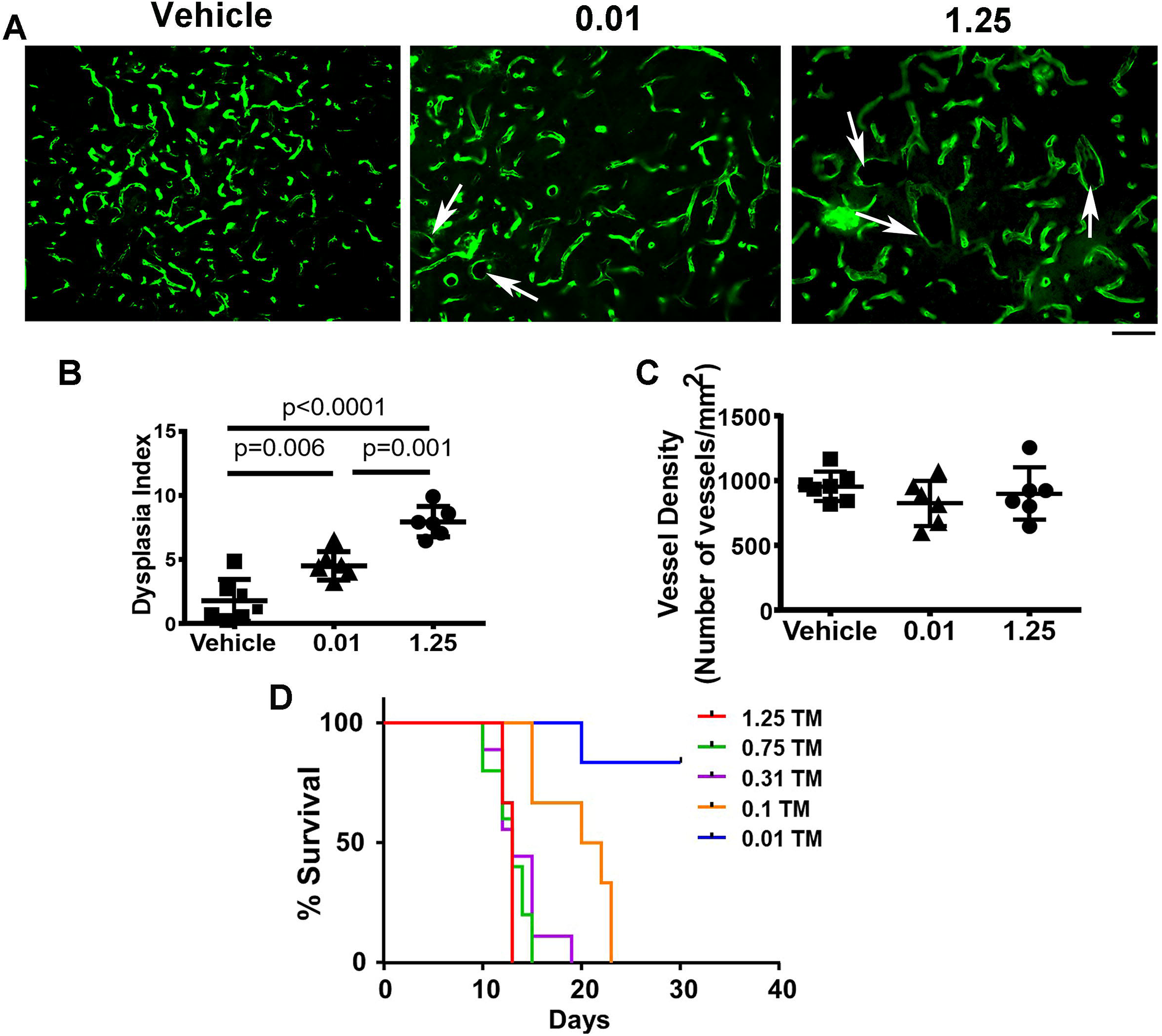
Increase of the TM dose increased the number of dysplastic vessels in the brain AVM lesions and mouse mortality. **A.** Representative images of sections collected from lectin (green) perfused brains. Arrows indicate dysplastic vessels. Scale bar: 80 μm. **B.** Quantification of dysplastic vessels. Dysplasia index: the number of vessels larger than 15 μm per 200 vessels. **C.** Quantification of vessel density. Vehicle: corn oil treated mice; 0.01 or 1.25: mice treated with 0.01 mg/25g or 1.25mg/25g TM. N=7 for Vehicle; N=6 for 0.01 and 1.25 groups. **D.** The survival curve. Mice were treated with one i.p. injection of 1.25 mg/25 g, 0.75 mg/25g, 0.31 mg/25g, and 0.1 mg/25g TM. N=6.

### Mutation of *Alk1* in BM-derived ECs alone is sufficient to cause AVM formation in the brains and AV shunts in the intestines

We and others have shown that mutation of AVM causative genes in ECs is essential for AVM development [6–8] and *Eng*-deficient BM transmits abnormal brain vascular phenotype to WT mice [11]. To test whether mutation of *Alk1* gene in BM-derived ECs is sufficient to induce AVMs in adult mice, we transplanted BM cells collected from *Pdgf*biCreER;*Alk1*^f/f^;Ai14^+/−^ mice to lethally irradiate 8-week-old WT mice. Brain angiogenesis was induced through an intra-brain injection of AAV-VEGF 4 weeks after the BM transplantation. Two weeks later, *Alk1* mutation in BM derived ECs was induced through one dose of i.p. injection of TM (2.5 mg/25g body weight, Fig. 6A).

**Fig. 6:**
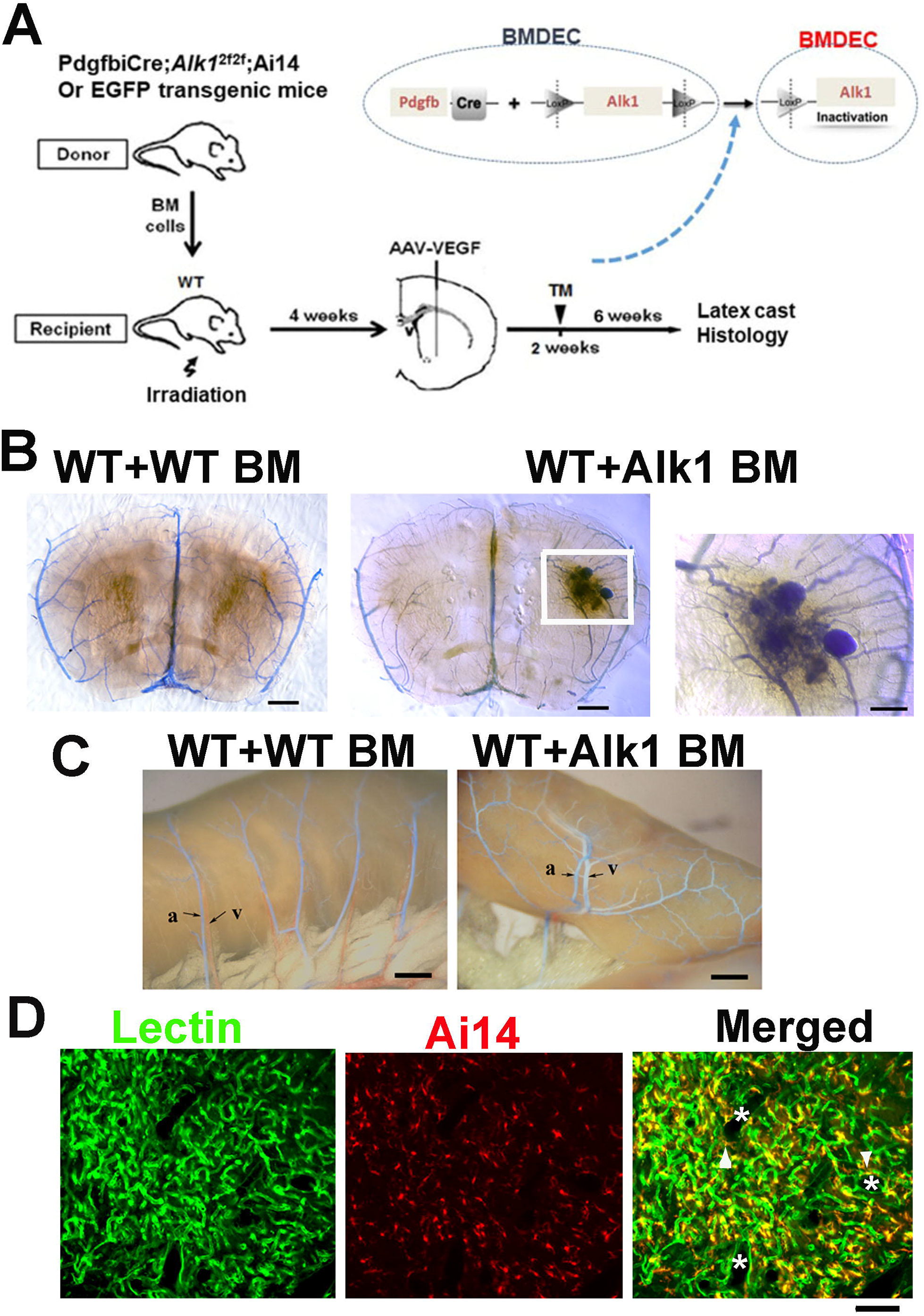
Mutation of *Alk1* in BM-derived ECs caused AVM development in the brain angiogenic region and AV shunts in the intestine. **A.** Study design for BM transplantation. *Pdgfb*iCre;*Alk1*^2f/2f^;Ai14^+/−^ or EGFP transgenic mice were used as BM donor. BM were transplanted to lethally irradiated WT mice. AAV-VEGF (2×10^9^ gcs) was injected to the brains of recipients 4 weeks after BM transplantation followed by TM (2.5 mg/25g body weight) treatment 2 weeks later to delete *Alk1* gene in BM-derived ECs. AVM phenotype was analyzed 6 weeks after TM treatment by latex cast and histology. **B**. Image of latex perfused brain sections. Left: A brain section of WT mice with WT BM. Middle: A brain section of WT mouse with *Pdgfbi*CreER;*Alk1*^2f/2f^;Ai14^+/−^ BM. AVM in the AAV-VEGF injected region is clearly shown. Right: Enlarged image of the white box in the middle picture shows the AVM nidus. Scale bars: 1 mm in Left and middle, 200 μm in Left. **C.** Latex casted intestinal vessels. Left image shows normal mesenteric vessels. Mesenteric veins (red) go parallel to the mesenteric arteries (blue latex perfused) and no latex dye was detected in the veins. Right image shows latex in the mesenteric veins, which indicates the presence of AV shunts in the intestine. a, artery; v, vein. Scale bars: 1 mm. **D.** Representative Images of brain sections collected from a lectin (green) perfused mouse with *Pdgf*iCreER;*Alk1*^f/f^;Ai14^+/−^ BM. Ai14 (red) was used as an indicator for Cre expressing BM-derived cells. Merged image shows Ai14^+^ cells are mostly co-localized with lectin positive ECs (green). *: dilated AVM vessels. Arrow heads indicate Ai14^+^ ECs on the abnormal vessels. Scale bars: 100 μm.

The recipients’ BM were fully reconstituted by donors’ BM 4 weeks after BM transplantation (Supplementary Fig. 4). Four of the 10 mice transplanted with *Pdgfb*-iCreER;*Alk1*^f/f^;Ai14^+/−^ BM developed different degrees of macroscopic level of AVM in the brain angiogenic regions and 5 of 9 mice with *Pdgfb*-iCreER;*Alk1*^f/f^;Ai14^+/−^ BM developed AV shunts in the intestines (Fig. 6B and C) after TM treatment and intra-brain injection of AAV-VEGF. Dilated AVM vessels were also detected on brain sections (Fig. 6D). There were more dysplastic vessels in brain angiogenic regions in mice transplanted with *Pdgfb*-iCreER;*Alk1*^f/f^;Ai14^+/−^ BM than mice transplanted with WT BM (P=0.006). Unlike *Pdgfb*iCreER;*Alk1*^f/f^;Ai14^+/−^ mice that died within 2 weeks after TM treatment due to intestinal bleeding and anemia [7], mice with *Pdgfb*iCreER;*Alk1*^f/f^;Ai14^+/−^ BM stayed viable beyond 6-week after TM treatment, suggesting that their intestinal vascular abnormality was less severe than *Pdgfb*iCreER;*Alk1*^f/f^;Ai14^+/−^ mice. These data indicate that BM derived ECs play a significant role in AVM development.

## DISCUSSION

In this paper, we showed that Alk1^−^ ECs undergo clonal expansion in brain AVM lesions. Equal numbers of Alk1^+^ and Alk1^−^ ECs were proliferating in the brain AVM lesions. Increase of TM dose increased the number of Alk1^−^ ECs in *Pdgfb*icre;*Alk1*^f/f^;Ai14^+/−^ mice and the AVM severity. Furthermore, we showed that deletion of *Alk1* gene in BM-derived ECs alone is sufficient to cause AVM formation.

Analysis of human brain and lung AVMs in HHT patients indicates that haploinsufficiency of HHT causative gene is not sufficient to cause lesion development [20]. Inactivation of the remaining WT allele appears to have a powerful effect, irrespective of the mechanism by which it is inactivated, e.g., loss of heterozygosity or loss of protein during inflammation [21, 22]. In mice, the loss of a single allele of HHT causative genes, such as *Eng* or *Alk1,* reproduces certain aspects of the human disease in animal models, primarily found in older animals [23, 24]. Brain AVM did not develop effectively in mice with haploinsufficiency of these genes. Loss of both alleles of any HHT-causative gene is embryonically lethal in mice [25, 26] and conditional (tissue/time-specific) homozygous deletion of *Eng* [21] or *Alk1* [27, 28] results in striking vascular malformations resembling the AVMs found in HHT. We found that the phenotypes of *Alk1* and *Eng* deletion mediated by the *R26*CreER promoter are different, presumably due to inefficiency of *Eng* gene deletion [6]. We also found that that both Eng^−^ and Eng^+^ ECs were present in the brain AVM lesions in the HHT1 mouse model [6].

Although it is known that mutation of AVM causative genes in ECs is essential for AVM development [6, 7, 9] and gene mutation in a small fraction of ECs can cause brain AVM [5, 10], currently, it is not clear how a small number of mutant ECs leads to brain AVM. It is also not clear if the mutant ECs in AVM are coming from a few parental mutant ECs. Using mice with TM controlled confetti transgene, we observed ECs expressing same confetti colors clustered in brain AVM lesions. This phenotype was not detected in brain angiogenic regions of WT mice, nor in the brains of mice with *Alk1* gene deletion alone. Our data suggest that Alk1 mutant ECs in brain AVMs undergo clonal expansion, which could partially explain why a fraction of mutant ECs can lead to AVM. Due to the high recombination rate of *Alk1* floxed allele (near 100%) and low recombination rate of confetti transgene in response to 2.5 mg/25g TM treatment, we could not analyze if non-mutant ECs are also clonally expanding in AVMs our model. We will consider using a different reporter line to address this question in future [29].

We showed previously that mutation of *Alk1* gene in ECs in the adult brain increased EC-proliferation in response to VEGF stimulation [7]. *Alk1* gene were deleted in almost all ECs in that study. The dysplastic vessels have more proliferating ECs than the surrounding normal capillaries. Jin et al. showed that deletion of *Eng* in ECs in neonates resulted in more proliferating Eng^−^ ECs than WT ECs in brain AVM lesions [15]. In our study, using an adult mouse model with mosaic pattern of Alk1^−^ and Alk1^+^ ECs, we found that equal number of Alk1^+^ and Alk1^−^ ECs were proliferating in the brain AVM lesions. Our data suggest that Alk1^−^ ECs promote growth of surrounding Alk1^+^ ECs.

Similarly, the clonal analysis of the mosaic retinal vasculature of *Eng* knockout mice revealed an equal proliferation rate of WT and Eng^−^ ECs in AVMs vessels in the retina, although more Eng^−^ ECs were proliferating in the vessels outside of AVMs, which suggests non-cell autonomous mechanisms contributing to the expansion of AVMs [15]. The roles of WT ECs in AVM pathogenesis remains to be elucidated.

In the study, we have also evaluated the relationship between the number of mutant ECs and AVM severity. By titrating the TM dose, we were able to reduce the number of Alk1^−^ ECs in *Pdgfb*icreER;*Alk1*^2f/2f^;Ai14^+/−^ mice. We found that mice with 1.25 mg/25g TM have more severe AVM phenotype than mice treated with 0.01TM. Therefore, strategies, such as transfusion of normal circulating ECs to patients, that can reduce the burden of mutant ECs could be developed into new therapies to potentially alleviate AVM phenotype severity.

Several studies, including our own, support a role of BM-derived cells in AVM formation [11]. After VEGF stimulation in the brain, WT mice with *Eng^+/−^* BM developed a similar degree of capillary dysplasia as mice with *Eng^+/−^* somatic and BM cells, suggesting that the loss of even one allele in BM-derived cells results in an abnormal vascular phenotype [11]. Similarly, in a converse experiment, *Eng^+/−^* mice transplanted with WT BM have fewer dysplastic vessels compared to *Eng^+/−^* mice with *Eng^+/−^* BM. These data suggest that BM-derived cells contribute to AVM development and that the tendency to form AVMs might be mitigated by transplantation of normal BM cells. However, it is not clear which cell-type in BM play a central role in AVM development. In this study, we transplanted BM isolated from *Pdgfb*icreER;*Alk1*^2f/2f^;Ai14^+/−^ mice to lethally irradiated WT mice. AVMs were induced after the BM of WT mice were reconstituted by *Pdgfb*icreER;*Alk1*^2f/2f^;Ai14^+/−^ BM. We showed in our previous study that BM cells enter brain angiogenic region after AAV-VEGF injection [11, 30]. Although majority of BM-derived cells differentiated into macrophages or leukocytes, about 7-10% of BM-derived cells differentiated into ECs and incorporated into vessels [11, 30]. Here, we induced *Alk1* gene deletion in BM-derived ECs two weeks after the induction of brain angiogenesis (Fig. 6A) through i.p. injection of TM. We detected AVMs in the brains and AV shunts in the intestines six week later. Our data indicate that BM-derived ECs is a the major player among all BM cells in AVM pathogenesis.

In summary, we showed that Alk1^−^ ECs undergo clonal expansion in the brain AVM lesions, which provides a partial explanation to the question of how a fraction of mutant ECs cause AVM. Equal numbers of Alk1^+^ and Alk1^−^ ECs were proliferating in brain AVMs. The number of Alk1^−^ ECs is positively correlated with AVM phenotype severity. We also showed that *Alk1* mutation in BM-derived ECs is sufficient to cause AVM in the brain and AV shunt in the intestine. Together, our data suggest that reduction of mutant EC-numbers can be a target for developing new therapies for alleviating the severity of AVM in HHT patients.

