## Supplemental data file for "*Alk1* mutant endothelial cells undergo clonal expansion in mouse brain arteriovenous malformations"

**Materials and Methods**

**Stereotactic Injection of AAV-VEGF into the brain**

To induce brain AVM, an adeno-associated viral vector expressing vascular endothelial growth factor (AAV-VEGF) {Shen, 2006 #18953} was stereotactically injected into the basal ganglia at the beginning of model induction (**Supplemental Fig. 1**). After induction of anesthesia by 4% isoflurane inhalation, mice were placed in a stereotactic frame with a holder (David Kopf Instruments, Tujunga, CA), and a burr hole was drilled in the pericranium, 2 mm lateral to the sagittal suture and 1 mm posterior to the coronal suture. Two μl viral suspension containing 2X10^9^ genome copies (gcs) of AAV-VEGF were stereotactically injected into the right basal ganglia, 3 mm underneath the brain surface, at a rate of 0.2 μl per minute using a Hamilton syringe. The needle was withdrawn after 10 minutes and the wound was closed with a 4-0 suture.

**Immunohistochemistry**

#### Brain samples were collected at the time indicated in Supplemental Fig. 1. For immunostaining, the brains were freshly frozen in dry ice and cut into 20-μm-thick coronal sections using a Leica CM1950 Cryostat (Leica Microsystems, Wetzlar, Germany). Sections were incubated at 4°C overnight with the following primary antibodies: anti-CD31 (1:100, Santa Cruz, Biotechnology, CA or 1:1500, R&D, Systems, Minneapolis, MN), anti-Alk1 (1:50, R&D, Systems), anti-Ki67 (1:50, eBioscience, San Diego, CA), anti-RFP (1:300 Takara Bio, Japan) and Anti-Erg (1:100, Abcam, Burlingame, CA). Alexa Fluor 594-conjugated (1:200), Alexa Fluor 488-conjugated (1:200, Invitrogen, Carlsbad, CA) or Alexa Fluor 647-conjugated (1:100) antibody was used as secondary antibody to visualize positive signals. Vectashield antifade mounting medium containing 4′, 6-diamidino-2-phenylindole (DAPI), Vector Laboratories, #H-1200, Burlingame, CA) was used to stain the cell nuclei and mount the slides.

**Quantification of Vessel Density, Dysplasia Index, Ai14^+^ cells, Alk1^-^/Alk+ cells and proliferating ECs**

For quantification of vascular density (number of vessels per mm^2^) and dysplasia index (the number of vessels larger than 15 µm per 200 vessels){Chen, 2013 #25301}, a group of mice were injected with a fluorescein labeled lectin (lycopersicon esculentum lectin, Vector Laboratories #FL-1171, Burlingame, CA) through jugular veins, after being anesthetized with isoflurane inhalation. Twenty minutes later, animals were perfused with PBS followed by 4% paraformaldehyde (PFA) intracardially. Brains and intestines were collected, incubated in 4% paraformaldehyde containing 20% sucrose for 2 days, and then frozen in dry ice, and sectioned into 20-μm-thick sections using a Leica CM1950 Cryostat (Leica Microsystems, Wetzlar, Germany).

Two sections per brain approximately 100 µm nostril or caudal to the injection site or intestines were selected for the quantifications. Brain sections were examined and imaged using a Keyence fluorescence microscopy under a 20X objective lens (Model BZ-9000, Keyence Corporation of America, Itasca, IL). A total of six images were taken from each brain, three from each brain section (to the right, to the left and below the injection site). The images were coded by a researcher who did not participate in the quantification. All quantifications were performed by at least two researchers who were blinded with the group assignment.

For quantification of Alk1^-^ and Alk1^+^ ECs and proliferating ECs, two sections per brain approximately 100 µm nostril or caudal to the injection site were selected from fresh frozen brain samples. Alk1^-^ and Alk1^+^ ECs were quantified on sections co-stained with anti-CD31 and anti-Alk1 antibodies. Proliferating ECs were quantified on sections co-stained with anti-Erg, anti-Alk1 and anti-Ki67 antibodies.

**Confocal Microscopy**

Confocal images were taken using a Zeiss 780 upright laser scanning confocal microscope (Carl Zeiss AG, Oberkochen, German) with a 34 detector array. Images were obtained with a 20X objective, with water immersion. Images are composed of a single 2 μm optical section and channels were overlaid in Image J 1.52.

**Latex Perfusion**

Mice were deep-anesthetized by isoflurane inhalation. The abdominal and thoracic cavities were opened. Both left and right atria were cut off. Blue latex dye (1 ml, Connecticut Valley Biological Supply Co. Burlington, NC) was injected into the left cardiac ventricle using a 21-gauge needle attached to a 5 ml syringe. The brains and intestines were harvested and fixed with 4% paraformaldehyde overnight. The arteriovenous shunts in the intestines were examined and imaged directly. The brains were dehydrated with methanol series and clarified with benzyl alcohol/benzyl benzoate (1:1 ratio). After clarification, the brains were cut coronally and imaged.

**Western blot**

Brain tissues (1 mm^3^) containing the vector-injection site were collected. Protein was extracted using a cell lysis buffer (Cell Signaling, Danvers, MA) supplemented with protease inhibitor cocktail (Sigma-Aldrich), 1mM PMSF (Cell Signaling) and quantified by the Bradford method (Bio-Rad, Hercules, CA). The protein concentration were determined using a microplate reader (Emax, Molecular Devices, Sunnyvale, CA). Protein samples 150-200 g were loaded into 4–20% Tris-Glycine gels (Bio-Rad, Berkeley, CA) and transferred onto nitrocellulose membranes (Bio-Rad). Immunoblotting were performed using primary antibodies specific to Alk1 (1:1000, Abcam) and Gapdh (1:1000, Abcam,). A donkey anti-rabbit IgG antibody was used as secondary antibody (Li-Cor, Lincoln, NE). Alk1 and Gapdh bands were detected by Li-Cor Quantitative western blot scanner and quantified using Li-Cor imaging software.

**Bone Marrow (BM) Transplantation**

BM transplantation was performed as previously described {Hao, 2008 #21718; Choi, 2013 #26231}. Briefly, BM cells were collected from the tibia and femurs of 8 to 10-week-old *Pdgfb*iCreER;*Alk1*^f/f^;Ai14^+/-^ mice or enhanced green fluorescent protein (EGFP) transgenic mice (the Jackson Laboratory, Bar harbor, ME) by flushing and aspiration with PBS containing 1% fetal bovine serum (FBS). Cells were centrifuged at 1,200 rpm for 10 minutes and resuspended in PBS at a concentration of 1X10^7^/ml. Two hundred μl of cell suspension (2X10^6^ cells) were injected into lethally irradiated (9.7 Gy, GC3000 Irradiator, MDS-Nordion, Ontario, Canada) wild type (WT) C57BL recipient mice (the Jackson Laboratory) via the tail vein. To facilitate the determination of the reconstitution rate, we transplanted BM cells collected from EGFP transgenic mice to WT control mice. Peripheral blood was collected 4 weeks after BM transplantation. Flow cytometric analysis (LSR II, BD Biosciences, San Jose, USA) was used to determine the nucleated cells in the recipients’ blood expressing GFP.

**Supplementary table 1.**

| **Penetrance of AV shunt in the intestines** | | |
| --- | --- | --- |
| **Vehicle** | **0.01** | **1.25** |
| **0% (0/6)** | **50% (3/6)** | **100% (6/6)** |

**Note**, 0.01and1.25: mice treated with 0.01 or 1.25 mg/25g TM.

N=6 per group

**Supplementary Figures.**

**Supplementary Fig. 1. Timelines for model induction and sample collection. A.** Timeline for mice that were used for analysis of clonal expansion. WT control mice were treated with TM vehicle (corn oil). **B.** Timeline for TM dose testing, gene expression analyses, and for vessel quantifications. TM 0.01/1.25: mice treated with 0.01 or 1.25 mg/25g TM. Day 22: samples were collected 22 days after intra-brain injection of AAV-VEGF from mice treated with 1.25 mg/25g; Day 28: samples were collected 28 days after intra-brain injection of AAV-VEGF from mice treated with 0.01 mg/25g

**Supplementary Fig. 2. Increase of TM dose increased the numbers for Ai14^+^ ECs in the intestines. A.** Representative images of intestinal sections. ECs were visualized by intravascular perfusion of fluorescent labeled lectin (green). Scale bar: 50 μm. **B**. Quantification of Ai14^+^ ECs in the intestines. N=5 for 1.25 group and N=4 for 0.01 group.

**Supplementary Fig. 3. Increase of TM dose increased the brain AVM severity and the penetrance and the number AV shunts in the intestines. A.** Brain vessels were casted with latex dye (blue). Scale bar: 200 μm. **B.** Representative images show AV shunts in the intestines. Scale bar: 1 mm. **C**. Quantification of AV shunts. 0.01 and 1.25: mice treated with 0.01 or 1.25 mg/25g TM. Vehicle: mice treated with corn oil.

The samples for 0.01 mg/25g TM treated mice were collected 6 days later than 1.25 TM treated mice (Please see Supplementary Fig. 1)

**Supplementary Fig. 4. Recipients’ bone marrow reconstituted by donor bone marrow.** Representative images of FACS analysis of GFP^+^ nucleated cells in the peripheral blood of wild-type mice transplanted with bone marrow of EGFP transgenic mice (WT/GFP) 4 weeks after the bone marrow transplantation. Wild-type mice (WT) or EGFP transgenic mice (GFP) were used as control.
