## Supplementary figures and images for "*Alk1* mutant endothelial cells undergo clonal expansion in mouse brain arteriovenous malformations"

### Sup Fig. 1

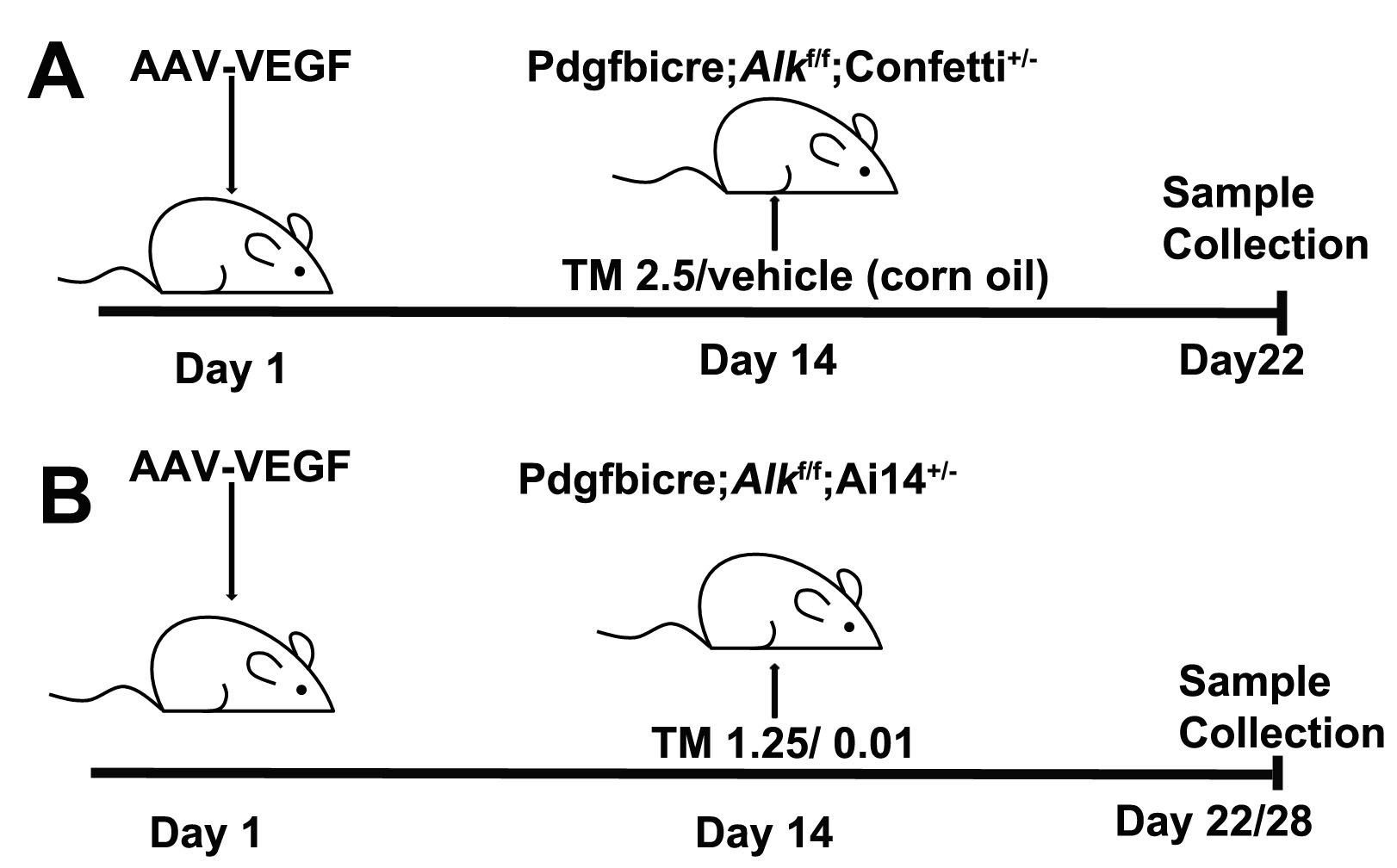

### Sup Fig. 2

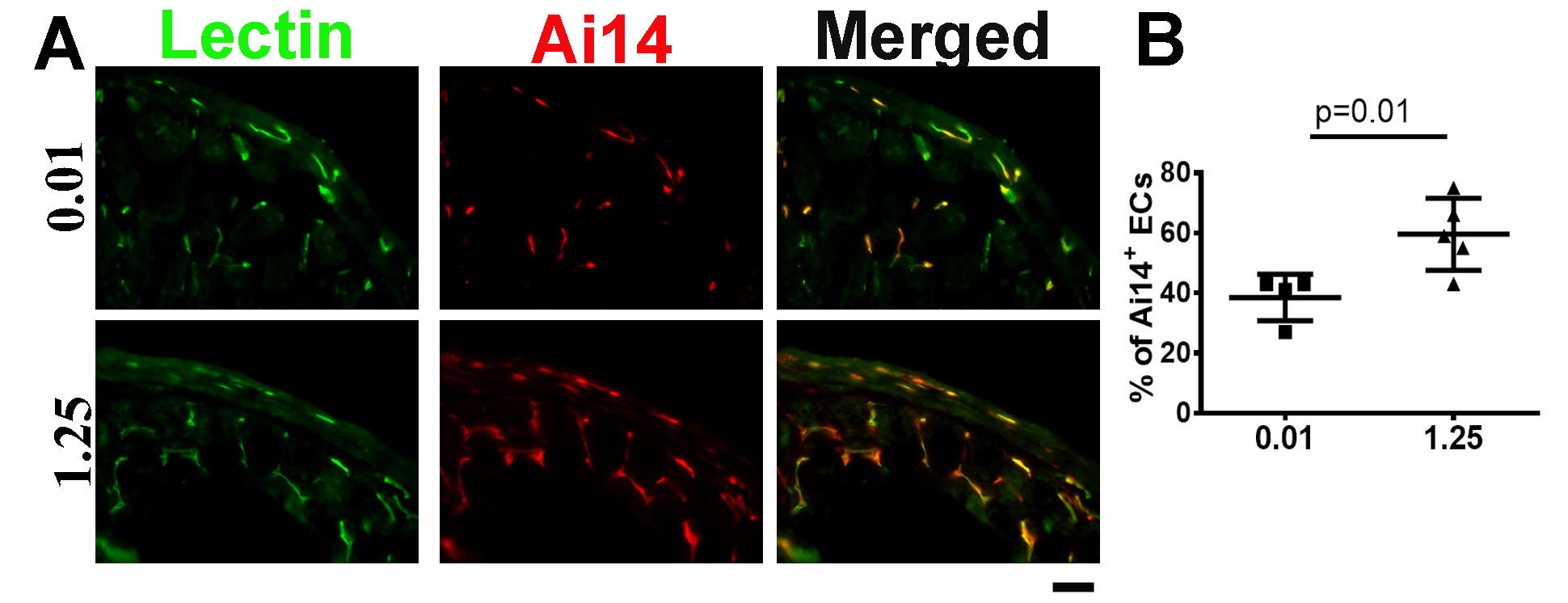

### Sup Fig. 3

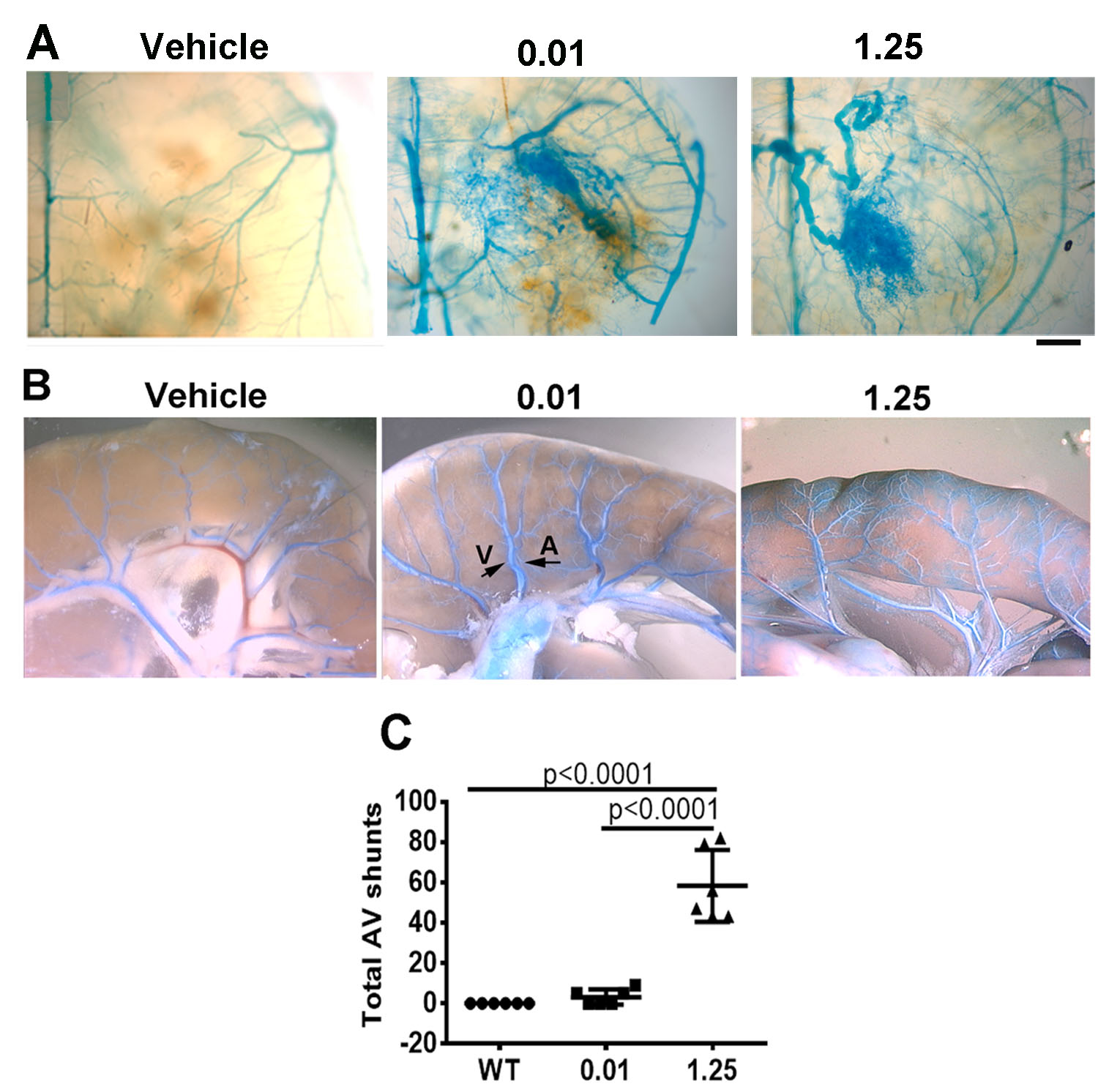

### Sup Fig. 4

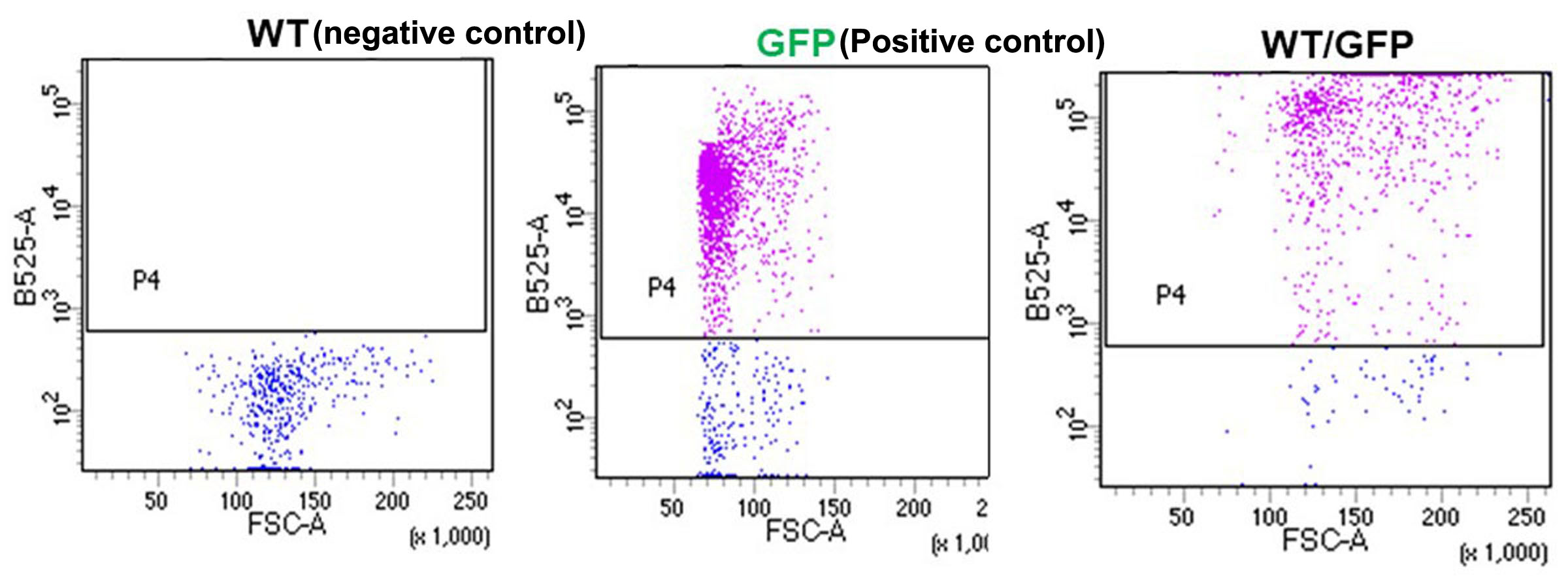
